# Shape Morphing Technique Can Accurately Predict Pelvic Bone Landmarks

**DOI:** 10.1101/2020.12.17.423253

**Authors:** Michal Kuchař, Petr Henyš, Pavel Rejtar, Petr Hájek

**Author notes:** Corresponding author *Email address:* (Petr Henyš).

## Abstract

Diffeomorphic shape registration allows for the seamless geometric alignment of shapes. In this study, we demonstrated the use of a registration algorithm to automatically seed anthropological landmarks on the CT images of the pelvis. We found a high correlation between manually and automatically seeded landmarks. The registration algorithm makes it possible to achieve a high degree of automation with the potential to reduce operator errors in the seeding of anthropological landmarks. The results of this study represent a promising step forward in effectively defining the anthropological measures of the human skeleton.

**Highlights:**

- The clinical CT scan is a feasible alternative to skeletal collections and body donor programs.
- Pelvic morphology is complex, sexually dimorphic and is proven to being a good demonstration model for the performance analysis of registration algorithm for automatic landmark seeding.
- The landmark seeding using registration algorithm can save time and effort in anthropological analysis.

## Introduction

In order to estimate the sex, body constitution or various anomalies of the individual from his/her skeleton, anthropologists typically rely on the nonmetric or metric analyses of the dry bone [1, 2]. More recently, the stereophotogrammetric method and medical imaging have been adopted [3]. In order to obtain reliable data, it is essential to work with a reasonably large set of specimens. The gold standard is the osteological database with personal data [4, 5, 6, 7]. Nowadays it is also possible to gain access to hospital databases and thus collect an equal or even larger set of data [8, 9] which is free from any postmortal changes. Virtual anthropometry is the method of choice in forensic cases [10], for the identification of victims of disasters [11] or in museum specimens that are susceptible to damage [12].

In recent years, we have been seeing rapid progress in the use of imaging techniques in forensic anthropology. Many studies have proven their compatibility with previous research on dry bone [13, 14] and found that CT scans are a promising source of reference data in contemporary forensic investigations [15, 16, 17]. It has been demonstrated that the accuracy of defining anthropological landmarks both manually and by use of CT scans have led to similar results between them [15, 18, 19, 20, 5]. Therefore the many methods that determine the sex of an individual, as well as the physical or biomechanical properties of a population that are already established and proven for skeletal material, could also be adopted for clinical CT data.

Regardless of the bony specimen‘s origin, its processing requires time and skill. We tried to reduce the time involved by adopting the technique of shape morphing for the mass analysis of anthropometric data. We adopted a non-linear registration algorithm which automatically computes the landmark positions from the ones that are pre–defined. The registration algorithm based on diffeomorphic mapping has been successfully used in brain analyses [21] but is rarely used in bone analysis [22, 23].

The study aimed to demonstrate the potential for shape registration in the automatisation of landmark seeding thus making data–gathering and evaluation easier in further studies, regardless of the researcher‘s experience. We created a set of virtual human pelvic bones and defined anatomical landmarks, which were automatically seeded by a proposed registration algorithm.

## Materials and Methods

### Dataset

Pelvic bone is well suited for our study because of its multifaceted morphology. Moreover, being the most sexually dimorphic skeletal element in the human body, it could further serve as a way of sex identification by using our proposed method. The basis for virtual modelling was the retrospective and anonymised DICOM files that were randomly taken from routine examinations in the Faculty hospital in Hradec Králové under ethical approval, 202010P08. The CT resolution of the data set was 1 × 1 × 1 mm (Siemens Definition AS+, Siemens Definition 128, 120-130 kV using CareDose, reconstruction kernel 80-90, bone algorithm). The inclusion criteria were: abdominal CT scans, bones without any trauma and an age range of 20 years or older. The sample population was equally balanced in terms of sex (100 males, 100 females), with the average age being 64 ± 13.5 years.

### The Segmentation of Bone Geometry

The pelvic bone geometries were obtained from CT scans with a semi– automatic segmentation algorithm (GraphCut, MITK-GEM, [24, 25]). On the downside, the algorithm may sometimes fail in finding the exact borders between the bones (sacral bone & pelvic bones, pelvic bones & the femur) that are fused via osteophytes. Therefore, in some cases, we had to manually correct the errors in the segmentation.

### Bone Registration

The purpose of image registration is to geometrically align the so called *moving image I* to the so called *fixed image J* by a suitable class of maps (see Figure 1). These maps transform each voxel **x** in the moving image *I*(**x**) to the corresponding voxel **y** in the fixed image *J* (**y**) by minimising a cost function that expresses the differences between *I*(**x**) and *J* (**y**) [26]. These transformations were computed by a well–known diffeomorphism method SYN in library ANTs with a modified intensity based criterion called the “demonslike metric” [27, 28]. The algorithm worked in the four–step resolution [100, 100, 50, 30] (the numbers in parentheses represent maximum optimisation iterations ^1^. In this study, we used those transformations to map anatomical landmarks from template shape onto a sample shape and vice versa^2^, see the convention in [26]. A suitable template bone must be created in such a way, that it minimises the anatomical discrepancies between the template bone and any sample it is morphed into. The template bone shape was iteratively estimated according to [27]. Once the template bone was obtained, all the samples in the dataset were morphed into the template bone shape. Each morphed bone sample was visually inspected for the presence of any errors.

**Figure 1:**
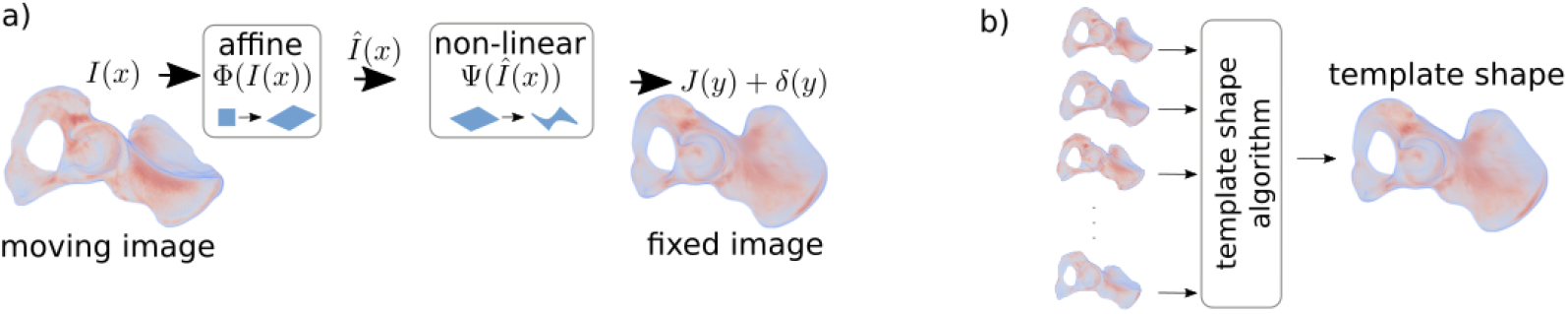
a) An illustration of the steps of the registration algorithm: the affine transform globally translates, rotates, scales and shears the moving image; the non–linear transform deforms (voxel–wise) the moving image in order to align the moving image with the fixed image. b) The fixed image is a template shape that is estimated from the dataset.

### Anthropological Measures

The template bone was set by a group of anthropometric reference land-marks *B*_1_, *B*_2_, …, *B*_19_ with the associated distances *M*_1_, *M*_2_, …, *M*_10_ (see Table 1 and Figure 2), by utilizing ParaView software [29]. We adopted the landmarks defined by Murail and Bruzek [30], both for their acceptance in the published literature [31] as well as for following the sex–specificity test. One additional landmark *B*_20_ was added to test the accuracy of the algorithm on the concave surfaces (on the bottom of the acetabular fossa).

**Table 1:** Definitions of the reference landmarks *B*.

|  |  |
| --- | --- |
| $B_1$ | Symphysis; the most superior and medial point on the pubic symphysis |
| $B_2$ | Anterior border of the acetabular rim at the level of the lunate surface |
| $B_3$ | The most lateral point on the acetabular rim |
| $B_4$ | A point on the medial margin of the pubic bone; at the level of $B_4$ |
| $B_5$ | The most inferior point of the os coxae |
| $B_6$ | The most superior point of the os coxae |
| $B_7$ | The posterior inferior iliac spine |
| $B_8$ | A point on the anterior margin of the great sciatic notch. |
| $B_9$ | The most anterior and inferior point on the ischial tuberosity |
| $B_{10}$ | The furthest point on the acetabular margin from $B_9$ |
| $B_{11}$ | Anterior superior iliac spine |
| $B_{12}$ | Posterior superior iliac spine |
| $B_{13}$ | Anterior inferior iliac spine |
| $B_{14}$ | The deepest point in the greater sciatic notch |
| $B_{15}$ | The contact point of the arcuate line and the auricular surface |
| $B_{16}$ | The midpoint of the anterior portion of the greater sciatic notch |
| $B_{17}$ | A point on the lateral border of the acetabulum; at the level of $B_{16}$ |
| $B_{18}$ | The most inferior point on the acetabular rim in the longitudinal axis of the ischium |
| $B_{19}$ | The most superior point on the acetabular rim on the longitudinal axis of the ischium |

**Figure 2:**
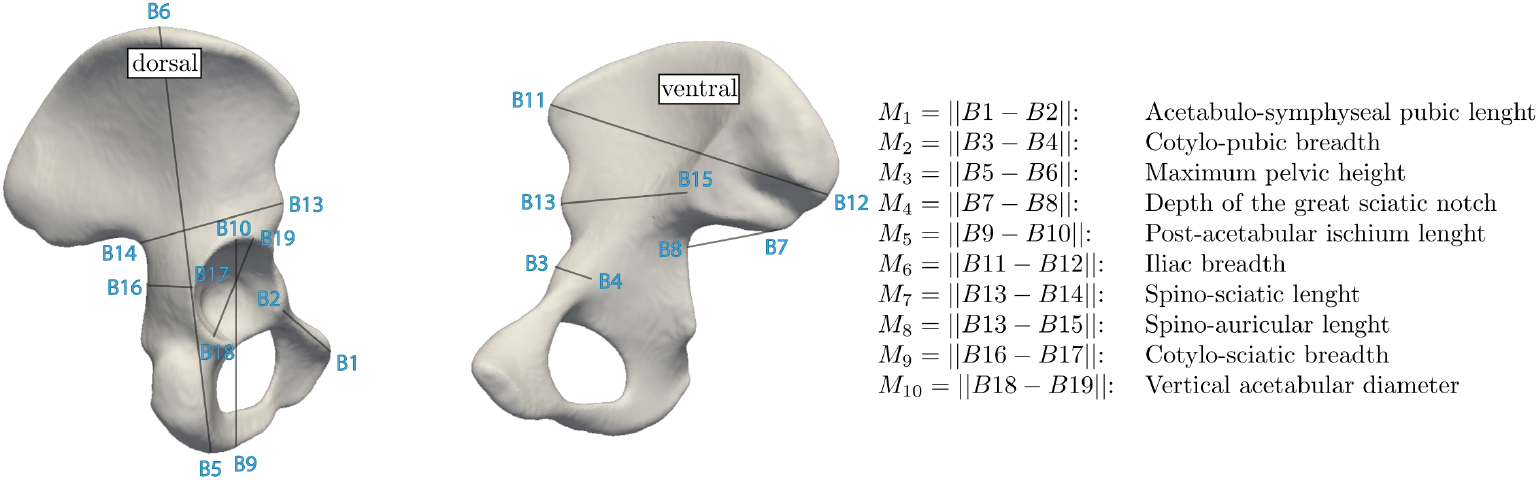
The estimated shape of the template bone with the reference landmarks *B* and distances *M*.

### A Comparison of Manually and Automatically Seeded Landmarks

In order to evaluate the accuracy of automatic the seeding algorithm, an operator manually seeded defined landmarks on 50 bones randomly selected from dataset.

### Intra–observer Error

We checked the consistency of manual seeding by analysing the intra– observer error in distances *M*. Fifty pelves were remeasured twice (test1 and test2) by a moderately experienced operator with a two week time window. The intra–observer technical error of measurement (TEM) and the percentages expressed relative rTEM were calculated. The resulting TEM index is a variable in anthropology that is used to express the margin of error and the quality of measurement. The mutual dependency of all tests is further expressed as the reliability coefficient R, that describes variance, which is free of measurement errors [32, 33, 34]:

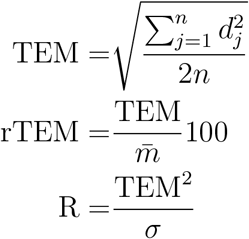

where *n* is the number of pelvis samples, 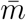 is the average distance value *M*, over the *n* samples, *σ* is the standard deviation over the *n* samples and *d*_*j*_ is the difference of *M* on the jth sample that is computed from the two measurements.

### The Distance Between Automatically and Manually Seeded Landmarks, B

To analyse the differences between both automatically and manually seeded landmarks, we computed the Euclidean distance

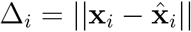

where **x**_*i*_ and 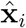 are the coordinates of the ith landmark *B*, that were obtained manually and automatically, respectively (see Figure 3). We analysed the distances on the samples from subsection Intra–observer Error. The statistical difference between landmarks *B*, measured at both repetitions was measured by the Mann Whitney test with a probability level of 95%.

### The differences Between Automatically and Manually Computed Distances, M

Relative differences between automatically and manually computed distances *M*, were analysed from samples of subsection Intra–observer Error, see Figure 4. The ith relative distance difference 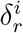 was computed as 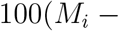 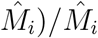. The statistical difference between distances *M*, measured at both repetitions was measured by the Mann Whitney test with a probability level of 95%.

**Figure 3:**
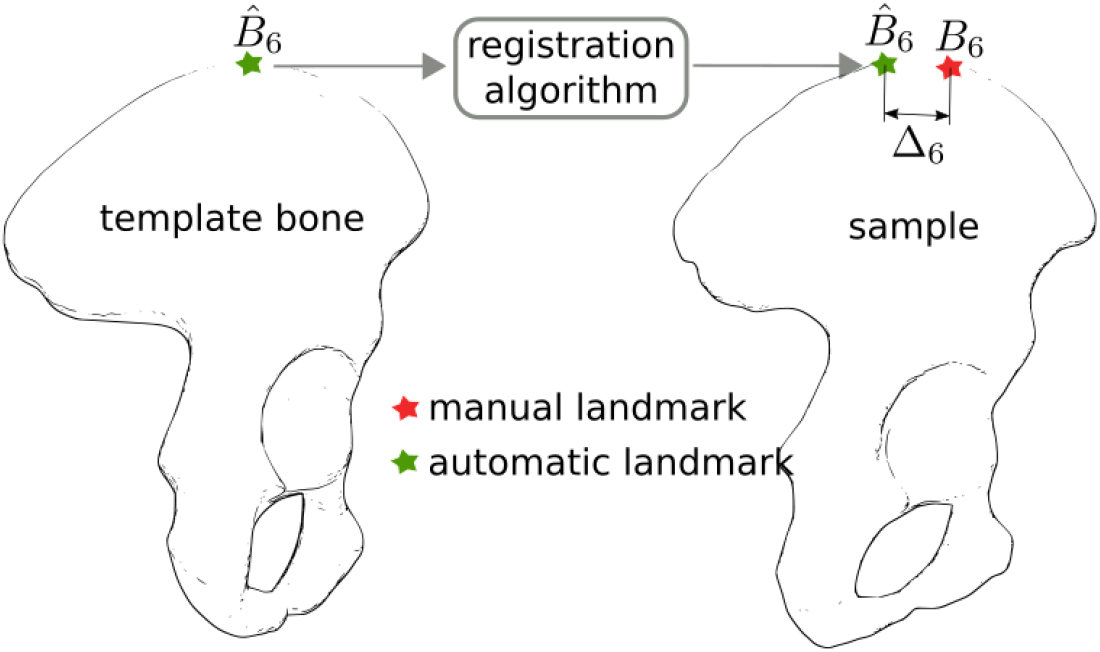
An example of measurement of distance between manually and automatically seeded landmark *B*_6_.

**Figure 4:**
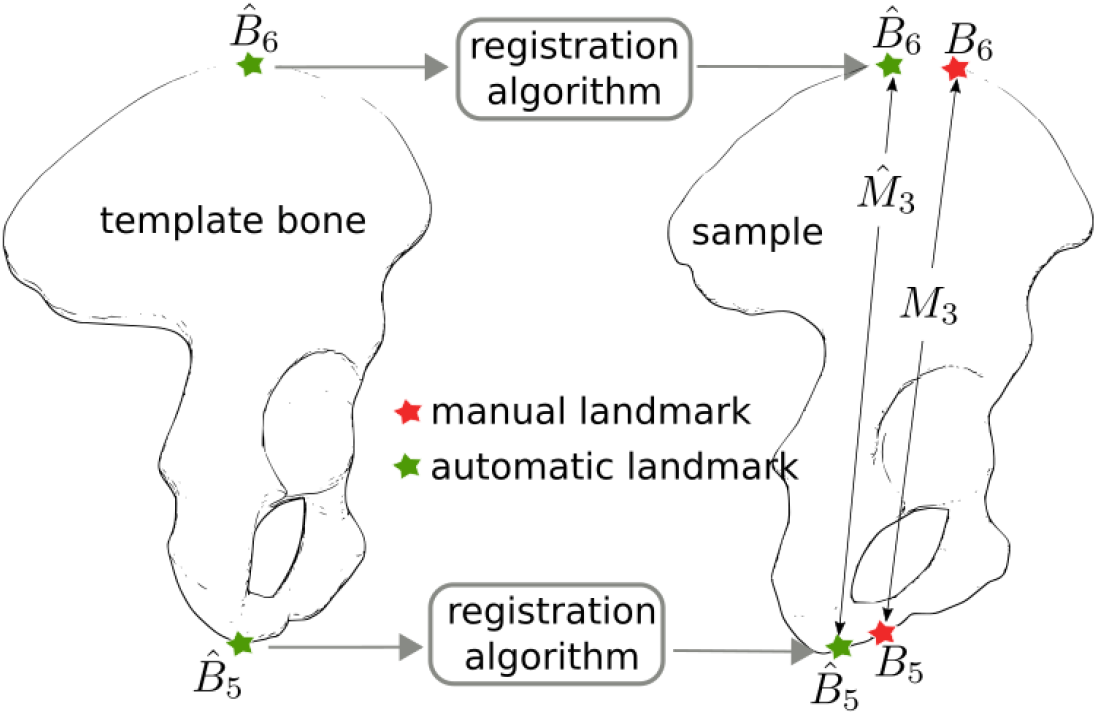
An example of the distance *M*_3_ computed from the manually seeded landmarks *B*_5_ and *B*_6_ and the distance 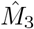 computed from automatically seeded landmarks 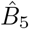 and 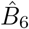.

### The Analysis of Clouds: the Back–Mapped Landmarks

The manually defined landmarks on the samples from subsection Intra– observer Error were mapped onto the bone template. The mapped landmarks form clouds around the reference landmarks. These landmark clouds have a certain shape, size and centroid (mean coordinates), which are used to analyze the accuracy of registration algorithms, see Figure 5. The centroids and confidence ellipsoids (eigenvalues of the covariance matrix) were estimated for the landmark clouds by the Quadratic Discriminant Classification Method (QDCM) [35]. By using the QDCM, we were able to estimate the probability that a given reference landmark belongs to the corresponding landmark cloud. The QDCM was trained by samples from subsection Intra–observer Error. The stratified KFold strategy with 3 folds and a train/test splitting at 70%/30%, was chosen in order to obtain the best accuracy [35]. The mean resultant train/test accuracy metrics were 92%±6.1%/90%±8.3%. Besides, we computed the distance Δ, between the centroids and the reference land-marks.

**Figure 5:**
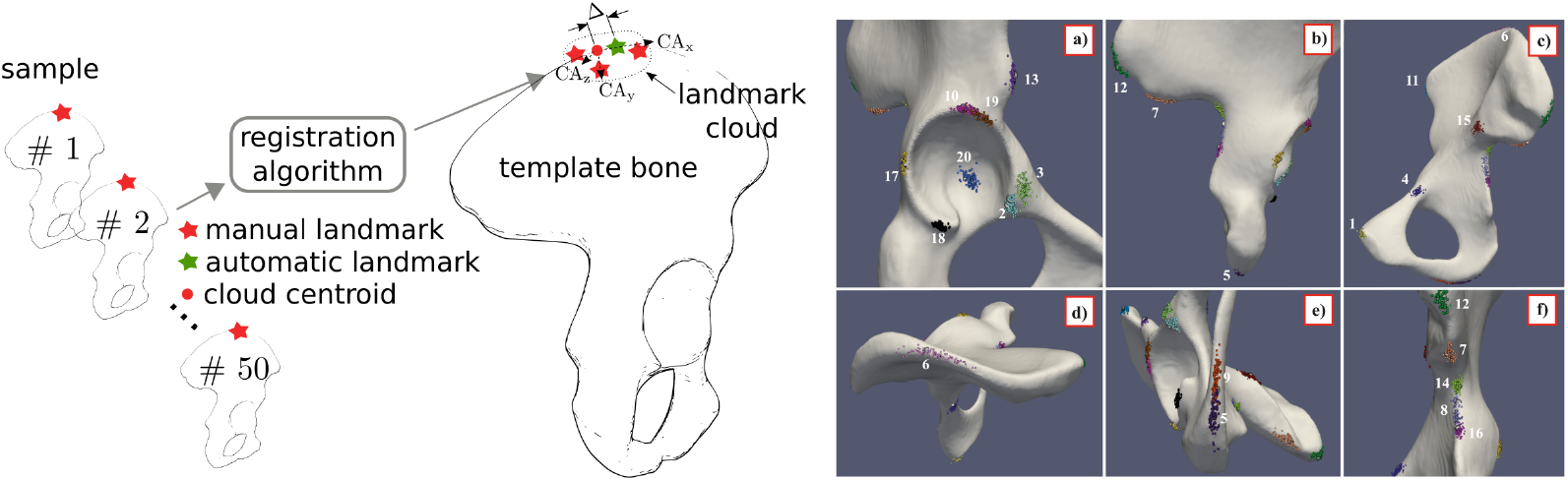
Use of registration algorithm for the mapping of manually seeded landmarks onto the bone template. The set [CA_*x*_ CA_*y*_ CA_*z*_] represents eigenvalues of a 95 % confidence ellipsoid. Individually coloured clouds are shown on various aspects of the pelvic bone [a), b), c), d), e), f)]. The numbers correspond to the landmark numbers in Figure 2.

### Sex Identification Using the DSP2 Method

We demonstrated the practical use and accuracy of the registration algorithm for sex identification. The input for the sex identification algorithm DSP2, were the distances from the subsection Intra–observer Error. DSP2 is based on a spreadsheet program (freely available at URL: http://projets.pacea.u-bordeaux.fr/logiciel/DSP2/dsp2.html) that gives the individual probability of being a male or female according to the linear discriminant analysis and posterior probabilities (see original publications [36, 30]). All ten distances *M*, served as an input to the application. The input was data consisted of 200 samples where the sex was known *a priori*.

## Results

### Observer Agreement

TEM values were in range of 0.60 for *M*_9_ and 1.55 for *M*_4_, see Table 2. The values of rTEM were mostly less than 2%, except for *M*_2_ and *M*_4_, which were 2.27 and 3.51 respectively and according to [37] are considered as being imprecise. The coefficient of reliability R, was between 0.94 and 0.99 and is defined as being high for all measurements. The TEM and rTEM were found as being relatively low [32].

**Table 2:** The technical and relative technical errors of manual measurements. The minimum and maximum values are in bold.

| | $M_1$ | $M_2$ | $M_3$ | $M_4$ | $M_5$ | $M_6$ | $M_7$ | $M_8$ | $M_9$ | $M_{10}$ |
| --- | --- | --- | --- | --- | --- | --- | --- | --- | --- | --- |
| TEM [mm] | 0.79 | 0.98 | 1.16 | **1.55** | 0.82 | 1.05 | 1.12 | 1.35 | **0.60** | 1.02 |
| rTEM [%] | 1.07 | 2.27 | 0.53 | **3.51** | 0.72 | **0.64** | 1.44 | 1.70 | 1.59 | 1.77 |
| R | 0.98 | 0.98 | **0.99** | **0.94** | 0.99 | 0.98 | 0.97 | 0.95 | 0.98 | 0.95 |

### The Distance Between the Automatically and Manually Seeded Landmarks, B

The largest average distance of 15.91 mm was found for landmark *B*_6_ while the smallest distance of 2.04 mm was found for landmark *B*_18_ in the test set 2, see Figure 6. There were no statistically significant differences between the repetitions of test1 and test2. The lowest value of p was 0.05 for landmark *B*_14_, while the highest value of 0.49 was found for landmark *B*_19_.

**Figure 6:**
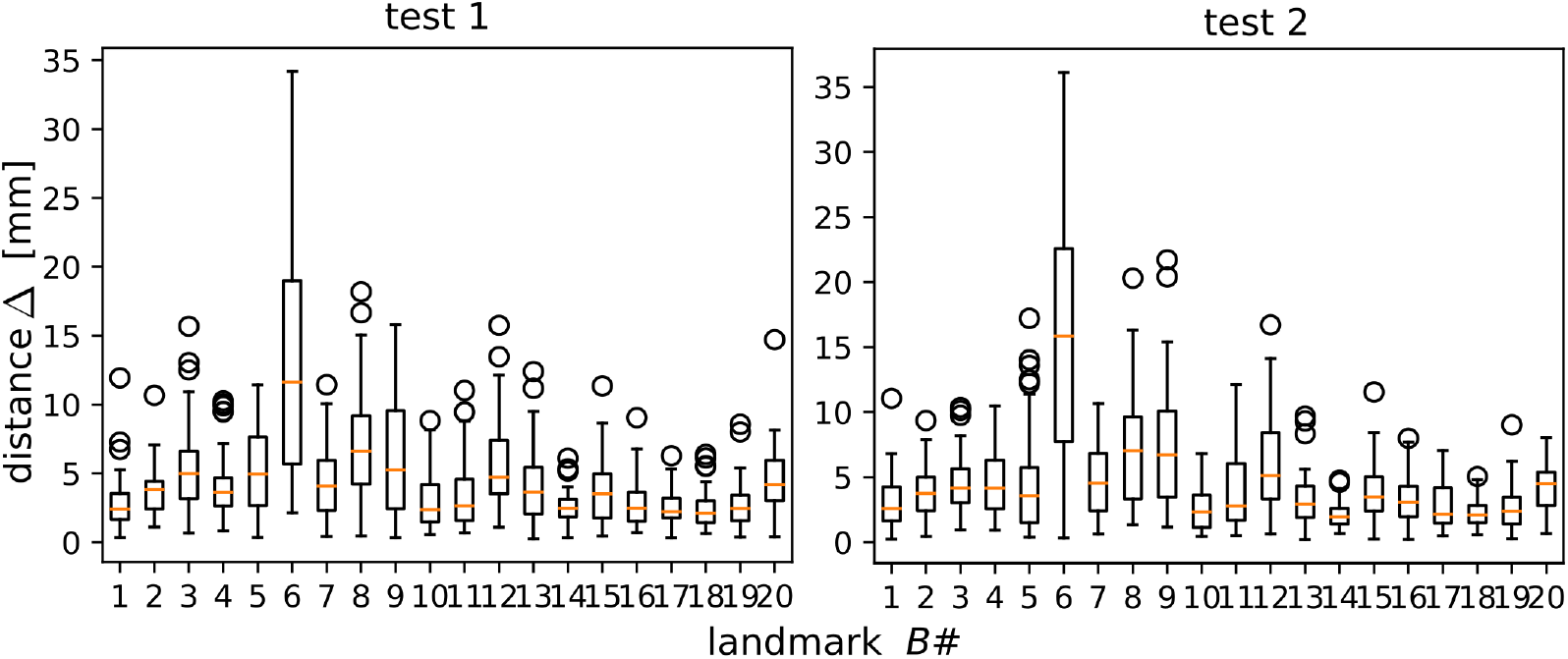
A boxplot showing the distance between the automatically and manually seeded landmarks for both repetitions.

### The Differences Between the Automatically and Manually Computed Distances, M

The largest average relative difference of −4.20% was found for distance *M*_4_ in the test set 2, see Figure 7. The average lowest relative difference of 0.01% was found for distance *M*_10_ in the test set 1, see Figure 7. There were no statistically significant differences between the repetitions of test1 and test2. The lowest value of p was 0.06 for the distance *M*_9_, while the highest value of 0.49 was found for distance *M*_2_.

**Figure 7:**
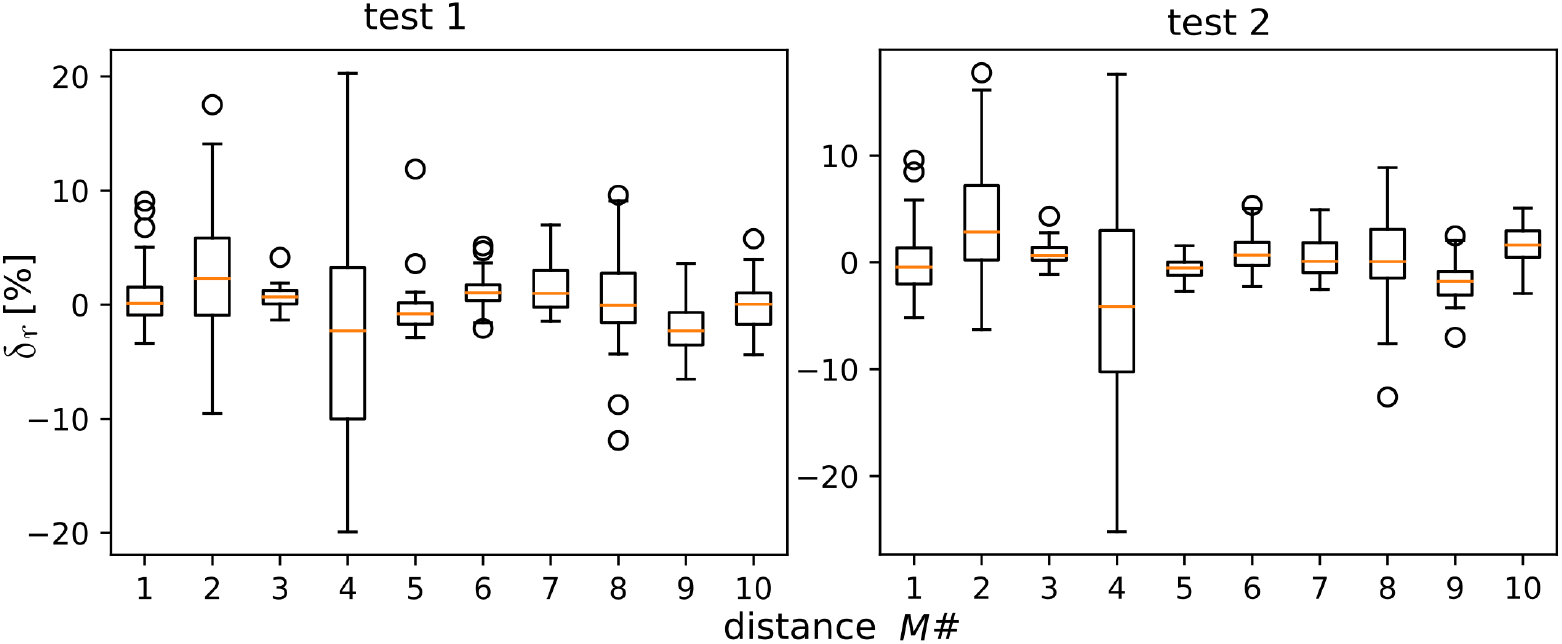
A boxplot showing the relative difference between automatically and manually seeded landmarks for both repetitions.

**Figure 8:**
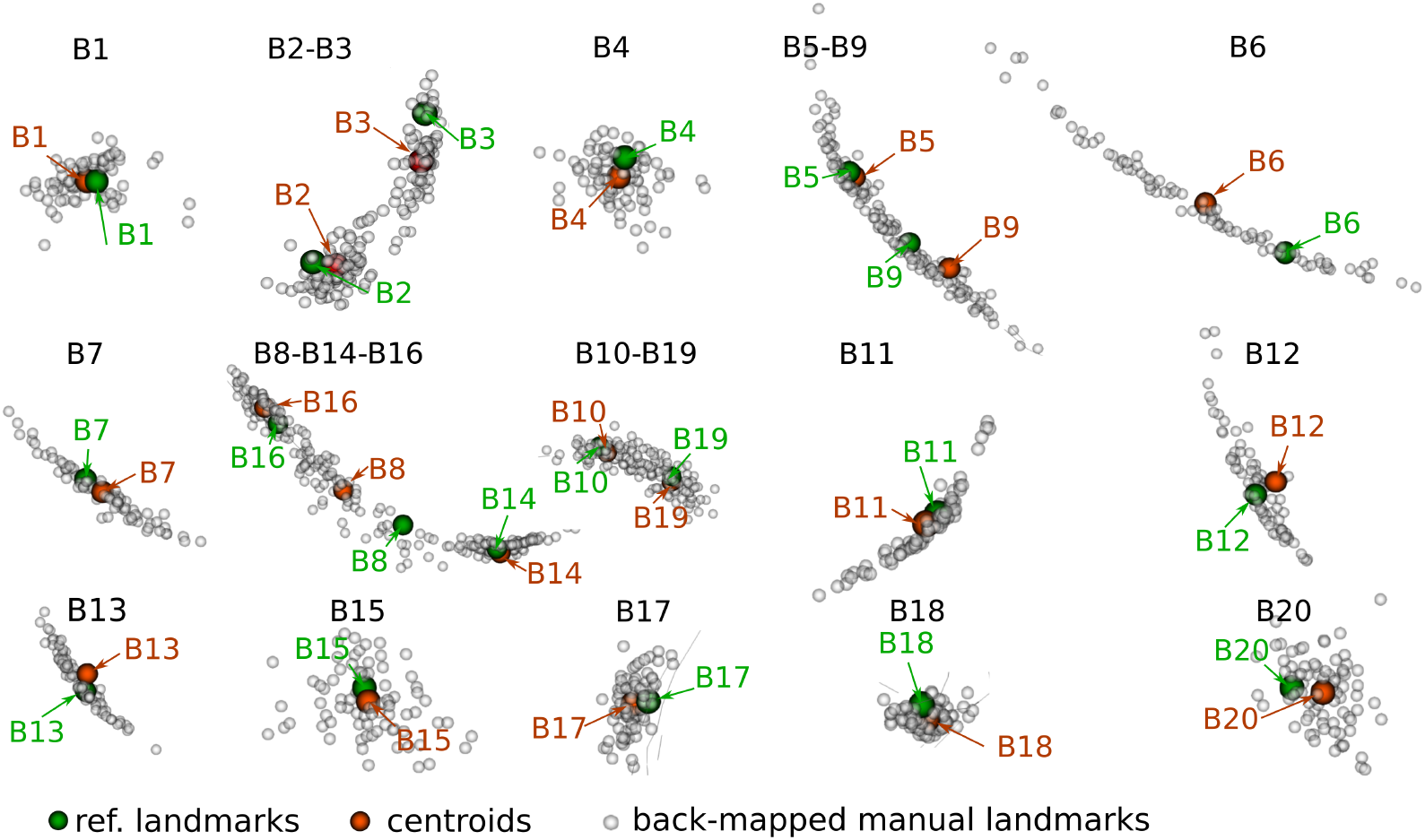
Clouds of manually seeded landmarks mapped onto the template bone.

### The Analysis of Clouds: Back–Projected Landmarks

The longest distance of 10 mm, was found between centroid *B*_6_ and reference landmark *B*_6_, while the shortest distance of 0.66 mm, was found between centroid *B*_19_ and reference point *B*_19_. The distances between the centroids and the reference landmarks are in Table 3. The probability that the reference landmark falls into a given landmark cloud was high (more than 99 %) for almost all landmarks. An exception was reference landmark *B*_9_, which fell into the landmark clouds of *B*_5_/*B*_9_ with a probability of 58.5%/41.5%, see Table 3. In addition, the highest length of confidence for axis x, was measured for point *B*_12_ with a value of 3.62 mm, while the lowest value, 0.677 mm was found for *B*_16_. The highest length of confidence for axis y, was found for landmark *B*_9_, with a value of 13.596 mm and the lowest was for landmark *B*_18_, with a value of 1.679 mm. The highest length of confidence for axis z, was found for point *B*_12_, with a value of 34.978 mm, while the lowest was for landmark *B*_18_, with a value of 3.978 mm.

**Table 3:** A comparison of the reference landmarks and centroids that are formed by a cloud of projected landmarks that were manually defined on a template bone;Δ[*mm*]) is the distance between the reference landmarks and centroids; [CA_*x*_ CA_*y*_ CA_*z*_] with the principal of a 95% confidence axes of an individual cloud. The minimum and maximum values are in bold.

| $B\#$ | $\Delta$ | $CA_x$ | $CA_y$ | $CA_z$ | $B\#$ | $\Delta$ | $CA_x$ | $CA_y$ | $CA_z$ |
| --- | --- | --- | --- | --- | --- | --- | --- | --- | --- |
| $B_1$ | 0.78 | 1.84 | 2.97 | 5.20 | $B_{11}$ | 1.20 | 1.40 | 2.58 | 7.11 |
| $B_2$ | 1.98 | 2.89 | 5.11 | 11.81 | $B_{12}$ | 2.12 | **3.62** | 8.83 | **34.97** |
| $B_3$ | 4.26 | 0.90 | 3.82 | 7.22 | $B_{13}$ | 1.63 | 2.68 | 6.24 | 16.38 |
| $B_4$ | 2.01 | 0.80 | 3.52 | 5.79 | $B_{14}$ | 0.79 | 2.70 | 3.99 | 16.36 |
| $B_5$ | 0.94 | 1.16 | 2.52 | 11.68 | $B_{15}$ | 1.19 | 0.69 | 4.62 | 7.58 |
| $B_6$ | **10.00** | 1.46 | 3.56 | 24.58 | $B_{16}$ | 2.03 | **0.67** | 2.26 | 4.05 |
| $B_7$ | 2.80 | 1.03 | 2.81 | 9.33 | $B_{17}$ | 1.27 | 0.79 | 2.02 | 4.82 |
| $B_8$ | 6.92 | 1.14 | 2.64 | 10.34 | $B_{18}$ | 1.41 | 0.90 | **1.67** | **3.97** |
| $B_9$ | 5.09 | 1.36 | **13.59** | 16.32 | $B_{19}$ | **0.66** | 1.05 | 2.06 | 6.84 |
| $B_{10}$ | 1.15 | 1.88 | 2.63 | 10.48 | $B_{20}$ | 2.50 | 1.03 | 3.79 | 6.02 |

### Sex identification Using the DSP2 Method

By using all ten distances *M*, 87% of males and 98% of females were successfully sexed. The sex was undetermined in 13 male and one female pelvic bone and wrongly assigned to one female. We excluded the distance *M*_4_ from sex identification as it was determined to being the most erroneous, see Figure 7. After excluding *M*_4_, an algorithm assigned 95% of the male pelves and 99% of the female pelves with a 100% accuracy in cases where the sex was assigned, see Table 4.

**Table 4:** Sex identification results by the DSP2 method (with a 95% posterior probability threshold). The variables on sexing accuracy and sex found to being indeterminable are expressed in %.

|  |  | Number of<br>distances M | Sexing<br>Accuracy | Sex<br>Undeterminable | Sexing<br>Error |
| --- | --- | --- | --- | --- | --- |
| Male | Original Setting | 10 | 87 | 13 | 0 |
|  | Corrected data | 9 | 95 | 5 | 0 |
| Female | Original Setting | 10 | 98 | 1 | 1 |
|  | Corrected Data | 9 | 99 | 1 | 0 |

## Discussion

Most of the average distances between the manually and automatically seeded landmarks were below 5 mm. Average distances above 5 mm were found for *B*_6_, *B*_8_ and *B*_9_. Landmarks *B*_6_ is defined from the distance *M*_3_ (see the definition in Figure 2), which means it is not directly dependent on bone geometry. Landmark *B*_6_ can be located almost anywhere in the middle third of the iliac crest. The landmark *B*_8_ should lie on a site, where the axis is inserted to the posterior inferior iliac spine just perpendicular to the anterior border of the greater sciatic notch (Figure 2). In this case, the operator’s result was superior to that of the computer’s result. This can be interpreted as an algorithm employing a similarity metric, which does not take into account any additional geometrical constraints.

The accuracy of automatic landmark seeding depends on the proper seeding of reference landmarks on template bone by an operator. Moreover, the identification of fine anatomical features on template bone can be more difficult because they can be partially smoothed out due to the method used for template bone construction [27]. This situation is typical for landmark *B*_9_, which relies on the location of the anteroinferior termination of the ischial tuberosity.

In our study, the TEM, rTEM and R values were relatively low and the mean differences between the automatic and manually measured distances were within millimetres which is comparable to similar publications [17, 14, 38, 39].

The sex identification results from the DSP2 method, with algorithmic computed input, proved to be very reliable. By leaving out distance *M*_4_ (which is dependent on *B*_8_), we achieved an improved sexing rate of 97% with a 100% accuracy, which is on par with the most relevant studies [14, 39, 38, 40].

The algorithm calculates a continuous spatial transformation, which means that any point on a bone sample has a unique counterpart on the template bone. In other words, we can potentially define landmarks anywhere on the bone [41]. This transformation makes it possible to interpret the difference in shapes in the deformation metric, which is considered as being intuitive and natural. This capability of the registration algorithm allows for shape analysis, which is usually performed by using the Principal Component Analysis (PCA) [42, 43, 44]. Unlike the PCA, the algorithm does not require a correlation matrix, which can be large and dense (in the case of CT data).

In our study, the algorithm took 10 minutes per sample (compiled on a Linux Ubuntu 18.04 LTS platform, GCC 7.4.0, Intel i7 (8 cores) CPU 2.10GHz, 16Gb RAM). This could be seen as a relatively long time, but the pipeline of registration is fully automated and stable, which is very convenient for the end–users. Once the registration step is done, the computing of landmark locations and distances over the whole dataset takes only a few seconds.

We are aware of some study limitations. Contrary to dry bone measurements, thin bone projections and bony plates could potentially be lost in the CT data due to an insufficient resolution and must be carefully reconstructed in order to obtain the same bone topology across the entire dataset. Furthermore, any articular surfaces that may be affected by entesophytes, which is common in the elderly, may reduce the accuracy of automatic landmark placement [45]. A lack of more observers should raise some caution regarding the interpretation of the intraobserver errors, however, a similar setup of the TEM method is proposed in [39, 46, 38].

## Conclusion

The anthropological landmarks must be seeded by an experienced operator, even when using CT data. Manually defining anthropological landmarks, especially for larger data sets, is inefficient and can introduce uncertainties depending on the operator’s focus on the work at hand as well as his/her level of experience. In this study, we introduced a method that allows us to automate the definition of anthropological landmarks based on a large amount of CT data. This method also makes it possible to potentially use data from bone digitization by laser scanning, which is a subject of further research. In summary, we

- introduced registration algorithm for automatic landmark seeding,
- extensively analysed the differences between manual and automatic landmark seeding, and
- showed the algorithm performance on the task of sex identification.

## Acknowledgements

This study was supported by a grant from the Grant Agency of Charles University SVV 260543/2020, Czech Republic. We would like to thank Dr Cyrus Rasti MD for his help in editing the English language content of this article.

## Footnotes

1 see the details in ANTs manual at URL: https://antspyx.readthedocs.io)

2 The points are transformed in the reverse direction unlike the images

## References

[1] W. Bass, Missouri archaeological society, Human osteology: a laboratory and field manual. 5th ed. Columbia (MO): Missouri Archaeological Society (2005).

[2] T. D., White, M. T., Black, P. A., Folkens, Human osteology, Academic press, 2011.

[3] K. Krishan, P. M., Chatterjee, T. Kanchan, S. Kaur, N. Baryah, R. Singh, A review of sex estimation techniques during examination of skeletal remains in forensic anthropology casework, Forensic science international 261 (2016) 165–e1.

[4] J. Bruzek, A method for visual determination of sex, using the human hip bone, American Journal of Physical Anthropology: The Official Publication of the American Association of Physical Anthropologists 117 (2002) 157–168.

[5] P. L., Walker, Greater sciatic notch morphology: sex, age, and population differences, American Journal of Physical Anthropology: The Official Publication of the American Association of Physical Anthropologists 127 (2005) 385–391.

[6] M. L., Patriquin, S. Loth, M. Steyn, Sexually dimorphic pelvic morphology in south african whites and blacks, Homo 53 (2003) 255–262.

[7] C. A., King, Osteometric assessment of 20th century skeletons from Thailand and Hong Kong, Universal–Publishers, 1997.

[8] W. S. Enns-Bray, H. Bahaloo, I. Fleps, Y. Pauchard, E. Taghizadeh, S. Sigurdsson, T. Aspelund, P. Büchler, T. B., Harris, V. Gudnason, S. J., Ferguson, H. Pálsson, B. Helgason, Biofidelic finite element models for accurately classifying hip fracture in a retrospective clinical study of elderly women from the ages reykjavik cohort., Bone 120 (2019) 25–37.

[9] D. Franklin, A. Cardini, A. Flavel, M. K., Marks, Morphometric analysis of pelvic sexual dimorphism in a contemporary western australian population, International journal of legal medicine 128 (2014) 861–872.

[10] M. A., Verhoff, F. Ramsthaler, J. Krähahn, U. Deml, R. J., Gille, S. Grabherr, M. J., Thali, K. Kreutz, Digital forensic osteology—possibilities in cooperation with the virtopsy^®^ project, Forensic science international 174 (2008) 152–156.

[11] M. Sidler, C. Jackowski, R. Dirnhofer, P. Vock, M. Thali, Use of multislice computed tomography in disaster victim identification—advantages and limitations, Forensic science international 169 (2007) 118–128.

[12] S. C., Kuzminsky, M. S., Gardiner, Three-dimensional laser scanning: potential uses for museum conservation and scientific research, Journal of Archaeological Science 39 (2012) 2744–2751.

[13] T. Chapman, P. Lefevre, P. Semal, F. Moiseev, V. Sholukha, S. Louryan, M. Rooze, S. V. S., Jan, Sex determination using the probabilistic sex diagnosis (dsp: Diagnose sexuelle probabiliste) tool in a virtual environment, Forensic science international 234 (2014) 189–e1.

[14] S. Mestekova, J. Bruzek, J. Veleminska, K. Chaumoitre, A test of the dsp sexing method on ct images from a modern french sample, Journal of forensic sciences 60 (2015) 1295–1299.

[15] K. E., Stull, M. L., Tise, Z. Ali, D. R., Fowler, Accuracy and reliability of measurements obtained from computed tomography 3d volume rendered images, Forensic science international 238 (2014) 133–140.

[16] K. L., Colman, A. E. van der Merwe, K. E., Stull, J. G., Dobbe, G. J., Streekstra, R. R. van Rijn, R.-J. Oostra, H. H. de Boer, The accuracy of 3d virtual bone models of the pelvis for morphological sex estimation, International journal of legal medicine 133 (2019) 1853–1860.

[17] K. L., Colman, H. H. de Boer, J. G., Dobbe, N. P., Liberton, K. E., Stull, M. van Eijnatten, G. J., Streekstra, R.-J. Oostra, R. R. van Rijn, A. E. van der Merwe, Virtual forensic anthropology: The accuracy of osteometric analysis of 3d bone models derived from clinical computed tomography (ct) scans, Forensic science international 304 (2019) 109963.

[18] D. Franklin, A. Cardini, A. Flavel, A. Kuliukas, M. K., Marks, R. Hart, C. Oxnard, P. O’Higgins, Concordance of traditional osteometric and volume-rendered msct interlandmark cranial measurements, International journal of legal medicine 127 (2013) 505–520.

[19] K. Aldridge, S. A., Boyadjiev, G. T., Capone, V. B., DeLeon, J. T., Richtsmeier, Precision and error of three-dimensional phenotypic measures acquired from 3dmd photogrammetric images, American journal of medical genetics Part A 138 (2005) 247–253.

[20] A. R., Klales, S. D., Ousley, J. M., Vollner, A revised method of sexing the human innominate using phenice’s nonmetric traits and statistical methods, American journal of physical anthropology 149 (2012) 104–114.

[21] N. J., Tustison, P. A., Cook, A. Klein, G. Song, S. R., Das, J. T., Duda, B. M., Kandel, N. van Strien, J. R., Stone, J. C., Gee, et al., Largescale evaluation of ants and freesurfer cortical thickness measurements, Neuroimage 99 (2014) 166–179.

[22] B. Dogdas, A. Chen, S. Mehta, T. Shah, B. Robinson, D. Xue, A. Gleason, L. D., Wise, R. Crawford, I. Pak, et al., Characterization of bone abnormalities from micro-ct images for evaluating drug toxicity in developmental and reproductive toxicology (dart) studies, in: 2015 IEEE 12th International Symposium on Biomedical Imaging (ISBI), IEEE, 2015, pp. 671–674.

[23] A. J., Ramme, K. Voss, J. Lesporis, M. S., Lendhey, T. R., Coughlin, E. J., Strauss, O. D., Kennedy, Automated bone segmentation and surface evaluation of a small animal model of post-traumatic osteoarthritis, Annals of biomedical engineering 45 (2017) 1227–1235.

[24] Y. Pauchard, T. Fitze, D. Browarnik, A. Eskandari, I. Pauchard, W. Enns-Bray, H. Pálsson, S. Sigurdsson, S. J., Ferguson, T. B., Harris, et al., Interactive graph-cut segmentation for fast creation of finite element models from clinical ct data for hip fracture prediction, Computer methods in biomechanics and biomedical engineering 19 (2016) 1693–1703.

[25] B. Helgason, S. Gilchrist, O. Ariza, P. Vogt, W. Enns-Bray, R. Widmer, T. Fitze, H. Pálsson, Y. Pauchard, P. Guy, et al., The influence of the modulus–density relationship and the material mapping method on the simulated mechanical response of the proximal femur in side-ways fall loading configuration, Medical engineering & physics 38 (2016) 679–689.

[26] B. B., Avants, P. T., Schoenemann, J. C., Gee, Lagrangian frame diffeomorphic image registration: Morphometric comparison of human and chimpanzee cortex, Medical image analysis 10 (2006) 397–412.

[27] B. B., Avants, N. J., Tustison, G. Song, P. A., Cook, A. Klein, J. C., Gee, A reproducible evaluation of ants similarity metric performance in brain image registration, Neuroimage 54 (2011) 2033–2044.

[28] B. B., Avants, N. Tustison, G. Song, Advanced normalization tools (ants), Insight j 2 (2009) 1–35.

[29] J. Ahrens, B. Geveci, C. Law, Paraview: An end-user tool for large data visualization, The visualization handbook 717 (2005).

[30] J. Bruzek, F. Santos, B. Dutailly, P. Murail, E. Cunha, Validation and reliability of the sex estimation of the human os coxae using freely available dsp2 software for bioarchaeology and forensic anthropology, American journal of physical anthropology 164 (2017) 440–449.

[31] G. Quatrehomme, I. Radoman, L. Nogueira, P. du Jardin, V. Alunni, Sex determination using the dsp (probabilistic sex diagnosis) method on the coxal bone: efficiency of method according to number of available variables, Forensic science international 272 (2017) 190–193.

[32] S. J., Ulijaszek, D. A., Kerr, Anthropometric measurement error and the assessment of nutritional status, British Journal of Nutrition 82 (1999) 165–177.

[33] R. Goto, N. M.-T. Cg, Precision of measurement as a component of human variation, Journal of physiological anthropology 26 (2007) 253–256.

[34] S. Carsley, P. C., Parkin, K. Tu, E. Pullenayegum, N. Persaud, J. L., Maguire, C. S., Birken, T. K., Collaboration, et al., Reliability of routinely collected anthropometric measurements in primary care, BMC medical research methodology 19 (2019) 84.

[35] F. Pedregosa, G. Varoquaux, A. Gramfort, V. Michel, B. Thirion, O. Grisel, M. Blondel, P. Prettenhofer, R. Weiss, V. Dubourg, J. Vanderplas, A. Passos, D. Cournapeau, M. Brucher, M. Perrot, E. Duchesnay, Scikit-learn: Machine learning in Python, Journal of Machine Learning Research 12 (2011) 2825–2830.

[36] P. Murail, J. Bruzek, F. Hoüet, E. Cunha, Dsp: a tool for probabilistic sex diagnosis using worldwide variability in hip-bone measurements, Bulletins et Mémoires de la Société d’Anthropologie de Paris (2005) 167–176.

[37] S. M., Weinberg, N. M., Scott, K. Neiswanger, M. L., Marazita, Intraobserver error associated with measurements of the hand, American Journal of Human Biology: The Official Journal of the Human Biology Association 17 (2005) 368–371.

[38] E. F., Kranioti, L. Št’ovíčková, M. A., Karell, J. Bruzek, Sex estimation of os coxae using dsp2 software: A validation study of a greek sample, Forensic science international 297 (2019) 371–e1.

[39] E. Vacca, G. Di Vella, Metric characterization of the human coxal bone on a recent italian sample and multivariate discriminant analysis to determine sex, Forensic science international 222 (2012) 401–e1.

[40] A. R., Paz, J. Banner, C. Villa, Validity of the probabilistic sex diagnosis method (dsp) on 3d ct-scans from modern danish population, La Revue de Médecine Légale 10 (2019) 43–49.

[41] M. Málková, J. Parus, I. Kolingerová, B. Beneš, An intuitive polygon morphing, The Visual Computer 26 (2010) 205–215.

[42] L. Savonnet, S. Duprey, S. V. S., Jan, X. Wang, Pelvis and femur shape prediction using principal component analysis for body model on seat comfort assessment. impact on the prediction of the used palpable anatomical landmarks as predictors, PloS one 14 (2019).

[43] S. Kai, T. Sato, Y. Koga, G. Omori, K. Kobayashi, M. Sakamoto, Y. Tanabe, Automatic construction of an anatomical coordinate system for three-dimensional bone models of the lower extremities–pelvis, femur, and tibia, Journal of biomechanics 47 (2014) 1229–1233.

[44] S. Schumann, L.-P. Nolte, G. Zheng, Comparison of partial least squares regression and principal component regression for pelvic shape prediction, Journal of biomechanics 46 (2013) 197–199.

[45] K. L., Colman, J. G., Dobbe, K. E., Stull, J. M., Ruijter, R.-J. Oostra, R. R. Van Rijn, A. E. Van der Merwe, H. H. De Boer, G. J., Streekstra, The geometrical precision of virtual bone models derived from clinical computed tomography data for forensic anthropology, International journal of legal medicine 131 (2017) 1155–1163.

[46] H. M., Karakas, A. Harma, B. Alicioglu, The subpubic angle in sex determination: anthropometric measurements and analyses on anatolian caucasians using multidetector computed tomography datasets, Journal of Forensic and Legal Medicine 20 (2013) 1004–1009.

